# A generalized machine-learning aided method for targeted identification of industrial enzymes from metagenome: a xylanase temperature dependence case study

**DOI:** 10.1101/826040

**Authors:** Mehdi Foroozandeh Shahraki, Kiana Farhadyar, Kaveh Kavousi, Mohammad Hadi Azarabad, Amin Boroomand, Shohreh Ariaeenejad, Ghasem Hosseini Salekdeh

**Affiliations:** Laboratory of Complex Biological Systems and Bioinformatics (CBB), Institute of Biochemistry and Biophysics (IBB), University of Tehran, Tehran, Iran; School of Natural Sciences, University of California Merced, Merced, California, United States of America; Department of Systems and Synthetic Biology, Agricultural Biotechnology Research Institute of Iran (ABRII), Agricultural Research Education and Extension Organization (AREO), Karaj, Iran; Department of Molecular Sciences, Macquarie University, Sydney, NSW, Australia

**Keywords:** machine learning, targeted identification, xylanase, metagenomics, thermal characterization

## Abstract

Growing industrial utilization of enzymes, and the increasing availability of metagenomic data highlights the demand for effective methods of targeted identification and verification of novel enzymes from various environmental microbiota. Xylanases are a class of enzymes with numerous industrial applications and are involved in the degradation of xylose, a component of lignocellulose. Optimum temperature of enzymes are essential factors to be considered when choosing appropriate biocatalysts for a particular purpose. Therefore, in-silico prediction of this attribute is a significant cost and time-effective step in the effort to characterize novel enzymes. The objective of this study was to develop a computational method to predict the thermal dependence of xylanases. This tool was then implemented for targeted screening of putative xylanases with specific thermal dependencies from metagenomic data and resulted in identification of three novel xylanases from sheep and cow rumen microbiota. Here we present TAXyl (Thermal Activity Prediction for Xylanase), a new sequence-based machine learning method that has been trained using a selected combination of various protein features. This random forest classifier discriminates non-thermophilic, thermophilic, and hyper-thermophilic xylanases. Model’s performance was evaluated through multiple iterations of six-fold cross-validations, and it exhibited a mean accuracy of ∼0.79. TAXyl is freely accessible as a web-service.

## 1. Introduction

Endo-1,4-beta-xylanase (EC 3.2.1.8) catalyzes the degradation of xylan, a component of hemicellulose, into xylooligosaccharides and D-xylose. Xylanases are currently being used in a broad spectrum of industries, such as pulp and paper, food, textiles, biofuel, animal feed, and beverage (D. Kumar et al., 2017). A considerable ratio of known xylanases is from glycoside hydrolases (GH) families 10 and 11 (Henrissat, 1991).

One of the essential attributes of enzymes is their optimum temperature, in which they exhibit their maximum relative activity. An enzyme’s optimum temperature is the indicator of its thermal activity (Liu, Wang, Zhang, Wang, & Chen, 2012). On the one hand, high temperature increases substrates’ solubility and bio-availability, accelerates molecular dynamics, and decreases the probability of microbial contamination significantly (S. Kumar, Dangi, Shukla, Baishya, & Khare, 2019; Vikash Kumar, Verma, Archana, & Satyanarayana, 2013). On the other hand, numerous biological processes are carried out at mild or cold temperatures (Collins, Gerday, & Feller, 2005). Therefore, various research studies have been focused on determining enzymes’ optimum temperature in order to identify appropriate novel biocatalysts for specific purposes (Li, Rabe, Nielsen, & Engqvist, 2019; Yan & Wu, 2012).

Unlike culture-dependent methods that are unable to cultivate more than 90% of microorganisms, culture-independent methods such as metagenomics enable the extended exploration of the natural diversity within an environmental sample (Schloss & Handelsman, 2003; Thomas, Gilbert, & Meyer, 2014). As the availability of metagenomic data is rapidly increasing, employing computational methods instead of wet-lab experiments in order to identify new enzymes with specific properties can effectively reduce the costs and make the process much faster. Microbial communities adjust to the requirements of their environment. For example, to enhance the digestion of plant matter and plant-derived complex polysaccharides, such as xylose and cellulose, which are the primary components of ruminant diet, rumen microbiome has acquired a rich hydrolyzing enzyme profile. (Stewart et al., 2019).

The strong correlation between the enzymes’ functional properties and its sequence suggest the importance of the primary structure of proteins in their attributes. Many studies have used sequence similarity-based methods in order to predict different properties of proteins. However, there are many cases where sequence similarity does not directly correlate with functional resemblance, as proteins with a nearly similar primary structure can sometimes depict unique properties or functional similarity can be observed among proteins with different amino acid sequences (Sadowski & Jones, 2009). Machine learning approaches have been successfully applied to predict various properties of proteins such as tertiary structure (Cheng, Tegge, & Baldi, 2008), function (Kulmanov, Khan, & Hoehndorf, 2018), localization (Almagro Armenteros, Sønderby, Sønderby, Nielsen, & Winther, 2017), thermal stability (Wu, Lee, Huang, Liu, & Horng, 2009), etc. These computational methods are capable of learning more complex relationships between the primary structure of proteins and their different properties (H. Bin Shen & Chou, 2006).

Numerous studies have presented *in-silico* methods for predicting enzymatic attributes. Discrimination between thermophilic and mesophilic proteins by using machine learning methods was the focus of a study by Gromiha and Suresh (Gromiha & Suresh, 2008). Similarly, Tang et al. used support vector machines (SVM) to develop a two-step method for discriminating thermophilic proteins (Tang et al., 2017) and amino acid compositions have been the basis of a statistical method for a similar task (Zhang, 2013). Pucci et al. presented a statistical approach to predict thermostability (Pucci, Dhanani, Dehouck, & Rooman, 2014), and Jia et al. designed a thermostability predictor tool (Jia, Yarlagadda, & Reed, 2015). AcalPred is another similar study that utilizes SVM to classify acidic and alkaline enzymes based on their primary structure (Lin, Chen, & Ding, 2013). In another research, Ariaeenejad et al. applied a regression model based on a pseudo amino acid composition (PAAC) (K.-C. Chou, 2009) to predict the optimum temperature and pH of xylanase in strains of Bacillus subtilis enzymes (Ariaeenejad et al., 2018). Genetic Algorithm-Artificial Neural Network (GA-ANN) have been employed for the optimization of xylanase production for industrial purposes (Vishal Kumar, Chhabra, & Shukla, 2017; Vishal Kumar, Kumar, Chhabra, & Shukla, 2019). In order to find features with the highest correlation with the thermostability of proteins, K-means clustering algorithm and a decision tree have been employed (Ebrahimi & Ebrahimie, 2010). Panja et al. found that the prevalence of smaller non-polar and hydrophobic amino-acids, as well as salt-bridges are some shared characteristics among most thermophilic proteins (Panja, Bandopadhyay, & Maiti, 2015).

The increasing use of new high throughput technologies can rapidly produce massive amounts of data, while the processes of getting access to the protein’s tertiary and quaternary structures are much slower. Therefore, an accurate and agile sequence-based approach is in demand to facilitate the targeted screening of high throughput data in order to find enzymes with the properties of interest. The objective of this study was to design and implement a multi-step method for the classification of the thermal activity of xylanases from glycoside hydrolases families 10 and 11 based on their optimum temperature. Since most of the available data in the literature belong to GH10 and GH11 families, we focused on the members of these two protein families. To the best of our knowledge, this is the first time that a combination of different protein descriptors is calculated, selected, and used to train a machine-learning classification model to make predictions on xylanase temperature dependence. The implemented tool has been successfully exploited for targeted isolation of three high performance xylanases from metagenomics samples. Moreover, we presented TAXyl (Thermal Activity prediction for Xylanase), a prediction web-server for the thermal activity of xylanases which was evaluated through multiple cross-validations as well as holdout test on unseen data.

## 2. Methods

### 2.1. Dataset preparation

A new dataset from GH families 10 and 11, which constitute a considerable ratio of known xylanases, had to be collected. Even though this makes the scope narrower, it can help the final estimation to be more accurate and reliable. The National Center for Biotechnology Information (NCBI) database was explored by searching for thermophilic or thermostable xylanases, and 254 results were found. After removing the records without the exact optimum temperature report, remaining sequences were divided into two groups of families 10 and 11. Afterward, the BRENDA (Jeske, Placzek, Schomburg, Chang, & Schomburg, 2019) and UniProt (Bateman, 2019) databases were explored for xylanases with reported optimum temperature, and newly collected data were added to the previous dataset.

Redundant or highly similar samples were removed using the CD-Hit (Huang, Niu, Gao, Fu, & Li, 2010), which clusters highly-homologous sequences and retains one representative sample from each cluster, with a 0.9 cut-off. In some protein sequences, a few amino acids are unknown. These residues are represented by the character “X.” Since the existence of such noise could potentially interfere with the learning process and feature extraction tools are designed for 20 amino acid residues, all unknown amino acids were removed from the sequences.

The final dataset consisted of 145 different xylanases from GH families 10 and 11 with optimum temperature ranging from 25°C to 95°C. These samples were labeled accordingly into three different classes: “Non-thermophilic” with optimum temperature below 50°C, “thermophilic” with the optimum temperature between 50°C and 75°C, and “hyper-thermophilic” with the optimum temperature above 75°C. Figure 1 shows the number of samples present in each thermal class and GH family.

**Figure 1.**
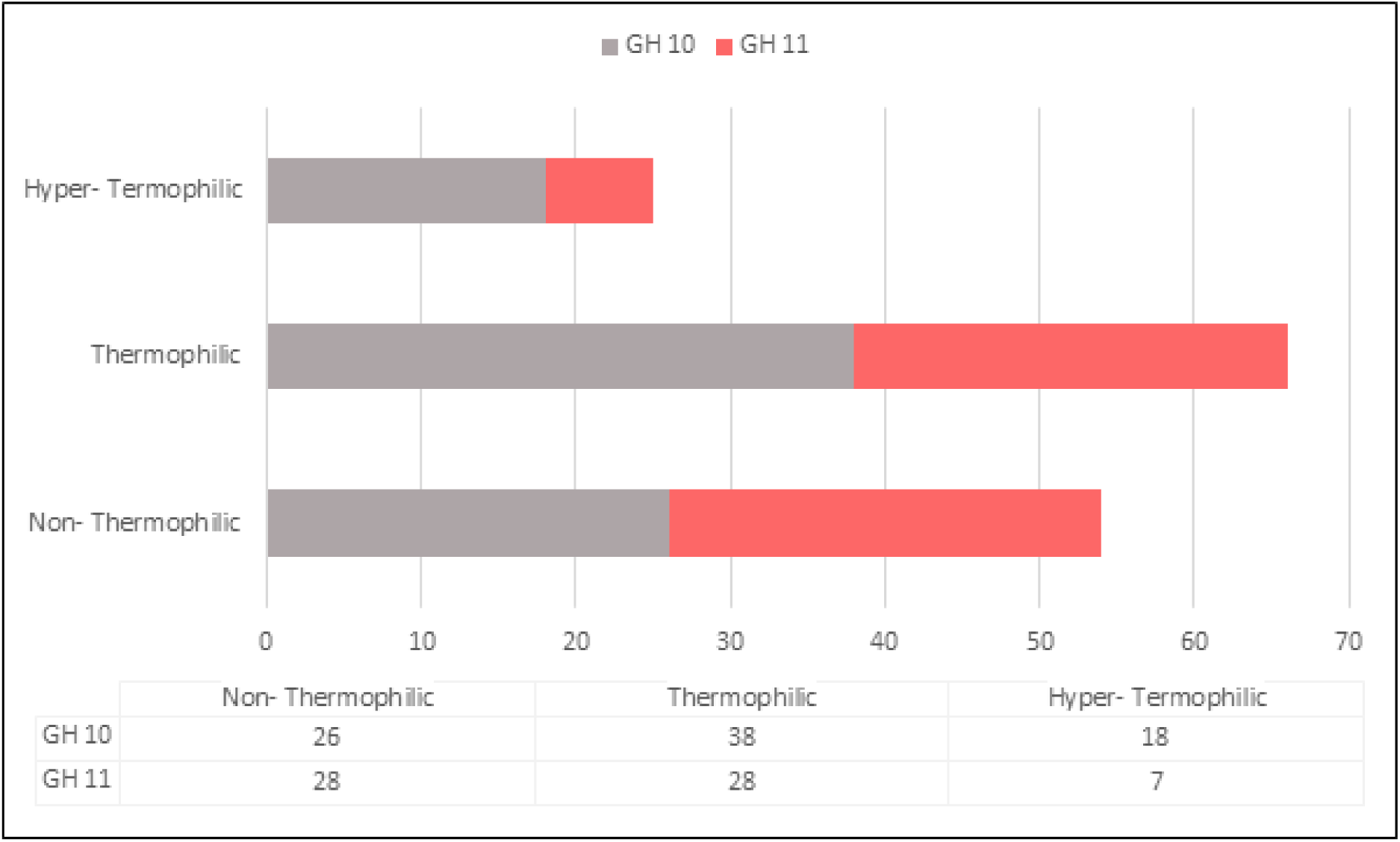
The dataset is divided into three categories. Samples in the dataset are from GH10 and GH11 families with optimum temperatures ranging from 25°C to 95°C. These enzyme samples are divided into three thermal activity classes. This figure illustrates the number of instances in each class and GH family.

### 2.2. Feature extraction and selection

Using an appropriate set of features is undoubtedly one of the most crucial steps in creating an efficient classifier. Because of the lack of precise evidence regarding the most related features to thermal activity, in this study, various protein descriptors were computed using the PyDPI python package (Cao et al., 2013).

The PyDPI-computed protein features are 15 descriptor types, which are from six main groups. Amino acid composition (AAC), dipeptide composition (2AAC) and tripeptide composition (3AAC), represent their fractions in the protein sequence. Unlike previous feature groups, pseudo amino acid composition (PAAC) and amphiphilic pseudo amino acid composition (APAAC), try to avoid missing the sequence-order information and reflect it in the composition data (H. B. Shen & Chou, 2008),(K. C. Chou, 2005). Conjoint triad features (CTF) cluster twenty amino acids into several classes based on their dipoles and the number of side chains (J. Shen et al., 2007). CTF considers the properties of an amino acid and its neighboring ones while regarding any amino acid triads as a unit. Other features are calculated by taking structural and physiochemical properties into account. Three autocorrelation descriptors, including normalized Moreau−Broto, Geary, and Moran autocorrelations, attempt to describe the amount of correlation among peptide or protein sequences. Composition, transition, distribution descriptors (CTD), sequence-order coupling number (SOCN), and quasi-sequence-order (QSO) describe the distribution pattern of amino acids along a protein sequence in terms of structural and physicochemical attributes.

As a result, a feature vector with 10,074 descriptors was calculated for each protein sequence. Many of these features were duplicates and were removed afterward. Due to different ranges of the descriptors, all raw values had to be re-scaled into same range.

All features do not contribute equally to the modeled response. Thus, a process of feature selection is necessary in order to find the most relevant descriptors for optimum temperature classification and to remove ones of less importance. Filter feature selection methods use a statistical measure to rank features based on their score. For this step, an F-test was applied as the filter method, and the best features were chosen using the SelectKBest method and the K-value was chosen to be 400 due to better performance.

### 2.3. Model selection and training

Numerous classification algorithms with different capabilities and weaknesses are currently available. In order to find the best methods for our problem, various classification algorithms were tested, including multi-layer perceptron (MLP), random forests, Gaussian Naïve Bayes, AdaBoost, and support vector machine (SVM) with radial basis kernel function (RBF).

Results of the model selection process suggested that random forest (RF) is the most suiting classification algorithm for this problem. RF is an ensemble machine learning method consisting of multiple decision trees which result in reduced variance and better generalization in comparison to single decision trees (Breiman, 2001). To build each tree of the RF, a bootstrap sample of the dataset was drawn randomly and the information gain was chosen as the splitting criterion.

In computational biology, RF is a popular method due to several advantages over other algorithms such as dealing with high-dimensional feature space, small number of samples, and complex data structures (Qi, 2012).

The above-mentioned pipeline required several hyper-parameter tuning steps, all of which were performed through grid searching. There are several approaches to detect the best combination of hyper-parameters for a machine learning model. The grid searching is a method which given multiple options for each hyper-parameter, tries every possible combination to build and evaluate the desired model. This method can thoroughly inspect the hyper-parameter-space in order to find the best configuration for the model to achieve the best performance.

### 2.4. Evaluation criteria

For the evaluation step, samples were randomly split into two sub-sample (90% as the training set and 10% as the holdout test set) ten times and each time fifteen iterations of 6 fold cross-validation (CV) were performed on the training set and after that, models were tested on the unseen sub-sample (holdout test set). This means that our classifiers were evaluated through 150 iterations of six-fold CV and they were tested on unseen data ten times to assure that the models are robust. In the six-fold CV step, the training set was again randomly split into six equal sub-sample, five of which were used as training sets and one sub-sample was then used for testing the models with different metrics. This validation process is executed 6 times leaving out one sub-sample each time for testing to ensure that all samples in the set were tested once. Figure 2 illustrates the evaluation process.

**Figure 2.**
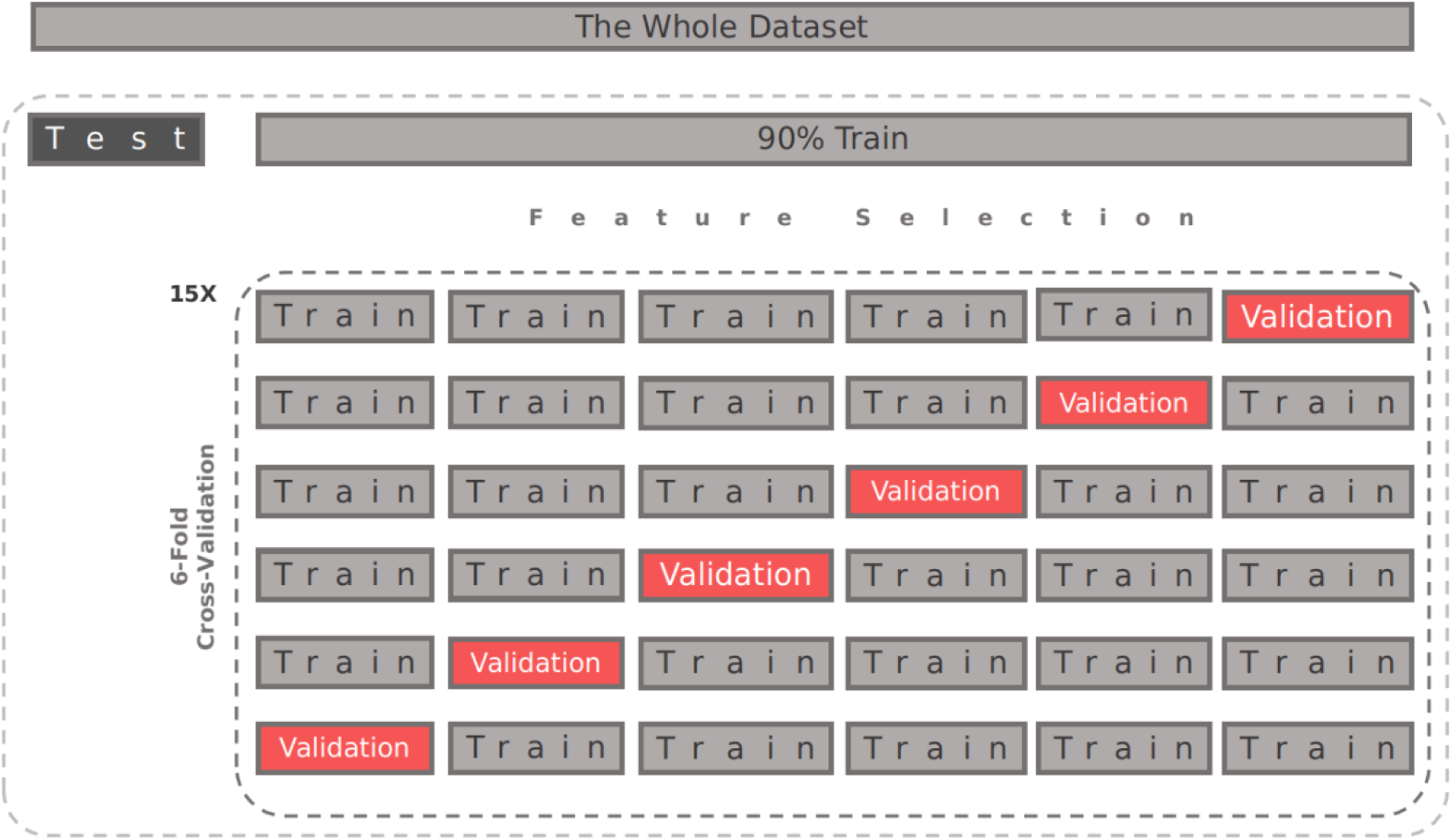
This figure demonstrates the evaluation process used to assess the predictive ability of the proposed model.

Since this is a multi-class classification problem, accuracy, macro-recall, macro-precision, the macro-f1 score, which are among the most commonly used metrics for classification tasks, were considered for evaluation of the model’s performance. These metrics were calculated using the following formulas:

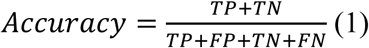

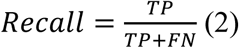

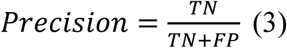

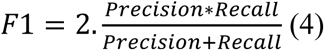

Here, TP (true positive) and TN (true negative) are positive and negative examples respectively that were correctly predicted. Similarly, FP (false positive) and FN (false negative) are positive and negative examples that were mistakenly classified. The “macro” prefix refers to averaging of each metric across the three different classes. The sci-kit learn Python package (0.21.2 version in Python 3.7.3) was used several times during selection, development, and evaluation of the classification model (Pedregosa FABIANPEDREGOSA et al., 2011). Figure 3 is the schematic diagram of the steps of development and evaluation of the current prediction model.

**Figure 3.**
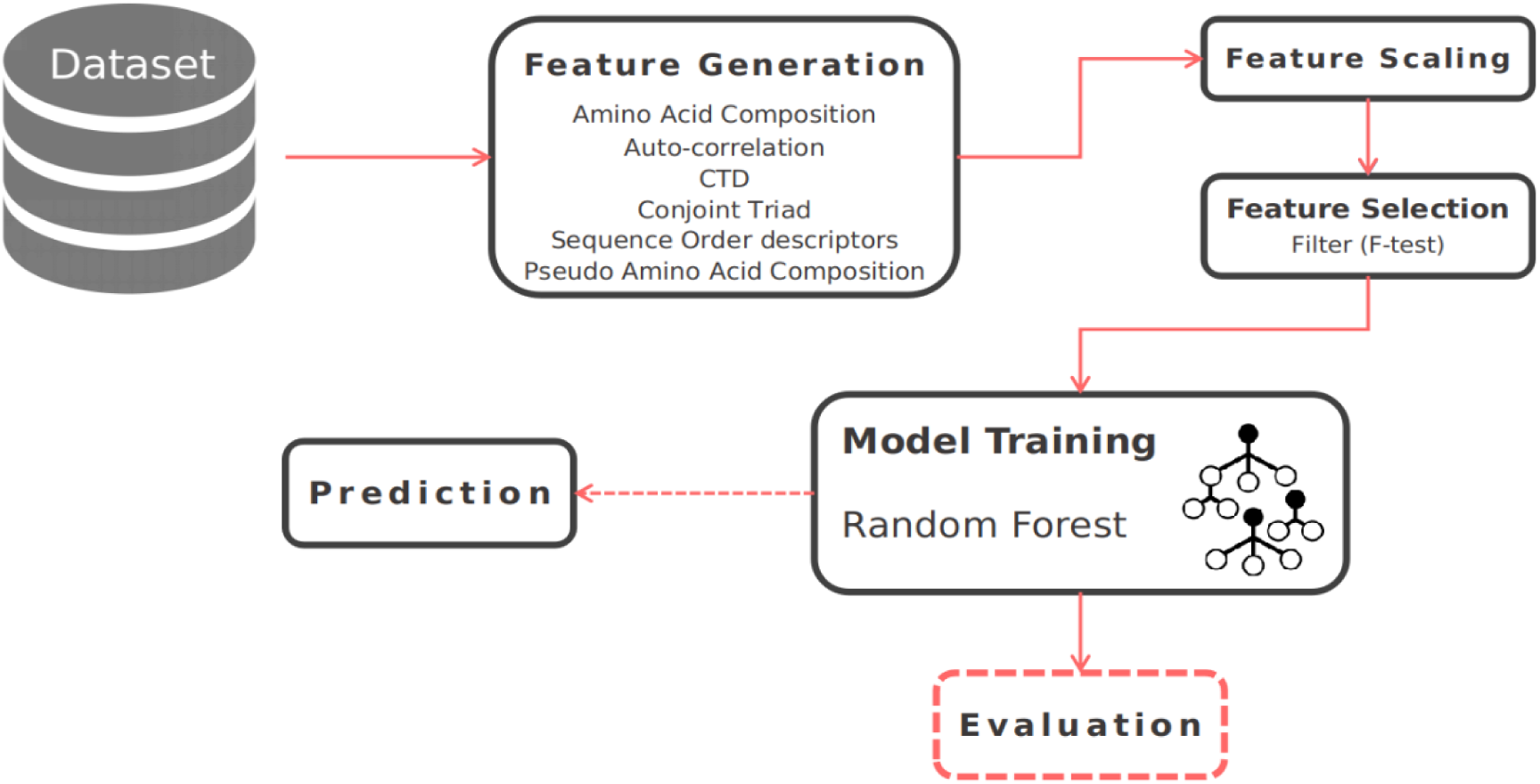
Schematic diagram of the workflow. The figure illustrates the steps for the development and evaluation of the proposed prediction model.

### 2.5. Cloning, expression, purification, and characterization of xylanases with predicted thermal activity

In order to verify TAXyl’s predictive capability, it was used to predict the thermal activity status of several metagenome-derived xylanase sequences all of which were subjected to different experimental analysis including determination of their temperature optima. Among putative xylanases from sheep and cow rumen metagenomic data, three candidates were nominated for cloning, expression and further experiments. Moreover, two xylanases from our previous studies were also used to validate the TAXyl’s ability in correct prediction of xylanases’ thermal activity.

In order to acquire xylanase genes, metagenomic DNA templates from sheep and cow rumen were used for polymerase chain reaction (PCR) amplification with two pairs of primers. To obtain the full-length coding genes, a specific primer set, the 5′- and 3′-ends of genes (Table 3) were designed for the PCR amplification. The resulting PCR products were detected on agarose gel 1.5% (w/v) and purified using the gel extraction kit (BioRon, Germany). Purified DNA fragments were cloned and digested into the pET28a. The resulting plasmids were then transformed into the E. coli DH5α for plasmid propagation followed by extraction and transformation of plasmid into E. coli BL21 (DE3) for xylanase expression. In the Luria-Bertoni (LB) medium, the recombinant strain pET28a was cultivated at the temperature of 37 °C. The cells were induced with isopropyl-β-thiogalactopyranoside (IPTG) for gene expression. The growing cells were then incubated at 19 °C for 19 h and finally were harvested by centrifugation at 4000×g for 30 min.

**Table 1.** A summary of different generated features from enzyme sequences that were used in this study.

| Feature Group | Number of Descriptors |
| --- | --- |
| Amino acid, Dipeptide and Tripeptide compositions | 8420 |
| Autocorrelations<br>(Moreau–Broto, Geary, and Moran) | 720 |
| Composition, Transition, Distribution | 147 |
| Conjoint triad | 512 |
| Quasi-Sequence-order and<br>Sequence-order coupling number | 190 |
| Pseudo Amino Acid Composition and<br>Amphiphilic Pseudo Amino Acid Composition | 85 |

**Table 2.**
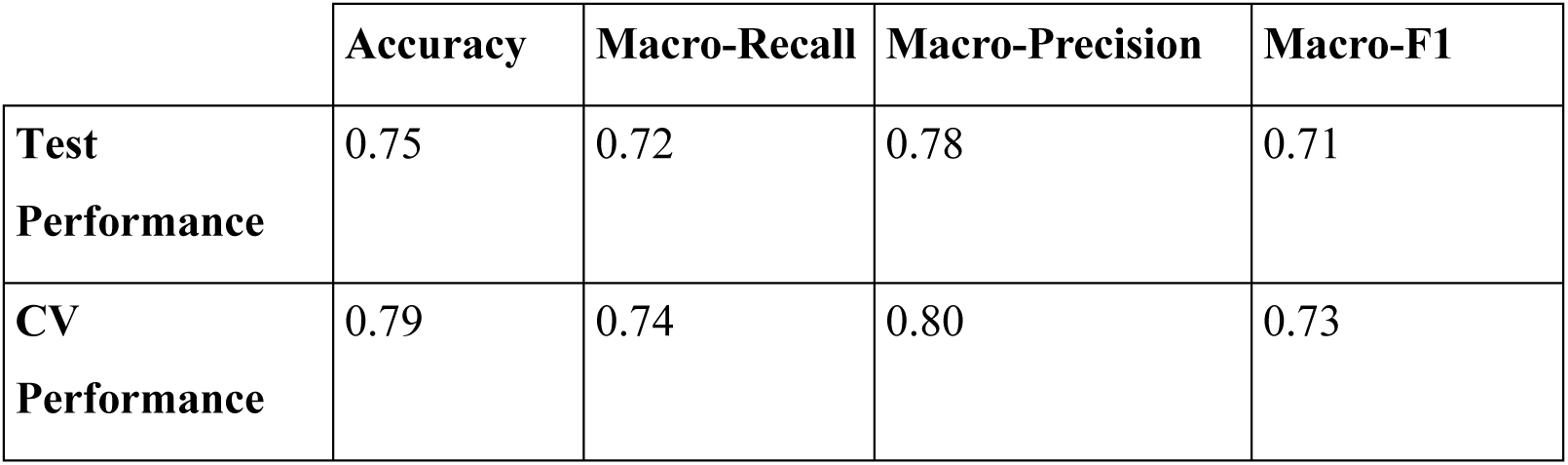
Comparison of cross-validation and holdout method test results. TAXyl’s performance during 100 iterations of 6-fold cross validations and five iterations of test (holdout) validations on previously unseen data.

**Table 3.**
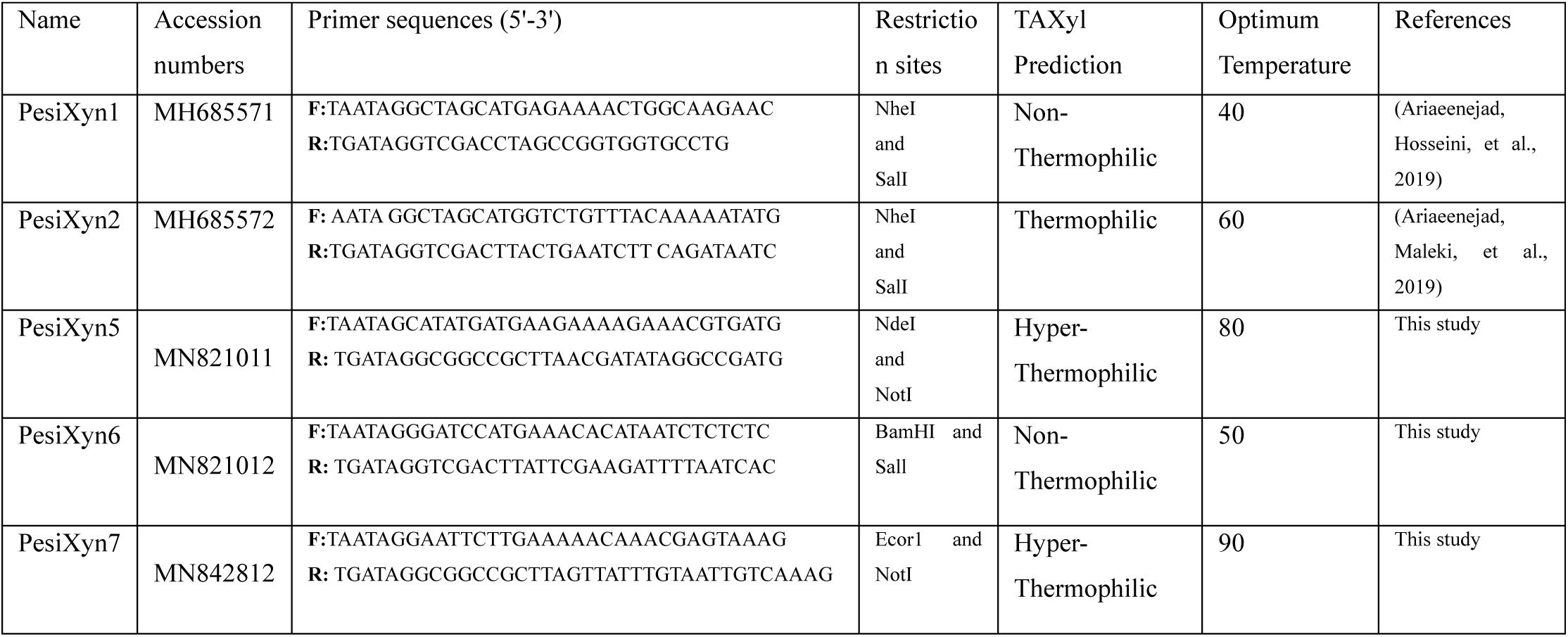
This table presents information regarding five metagenome-derived xylanases. Except for PersiXyn1 and PersiXyn2 which were from previous studies and were used only for validation purposes, TAXyl’s predictions were made prior to cloning and production of three remaining enzymes and facilitated the targeted isolation of these xylanases.

By utilizing Ni-NTA Fast Start Kit (Qiagen, Hilden, Germany), N-terminal Histidine-tagged recombinant protein was purified and evaluated by sodium dodecyl sulfate-polyacrylamide gel electrophoresis (SDS-PAGE).

Protein concentrations were determined through the Bradford method using the bovine serum albumin as the standard. By measuring the optical density (OD) of chromatography eluent at 280nm, the protein concentration was estimated.

To determine the optimum temperature of enzyme activity, enzyme solution in 10 mM phosphate buffer (pH 8) with substrate was incubated in the different temperature (30–90 °C) for 20 min and DNS was used to measure activity (Ariaeenejad, Hosseini, et al., 2019; Ariaeenejad, Maleki, et al., 2019). For reporting purposes, relative activity of each enzyme is considered as percentage of the highest demonstrated activity.

## 3. Results

### 3.1. Feature Extraction and Selection

Generated protein descriptors encapsulate various molecular and sequential information with different degrees of relevance to the enzyme’s thermal activity. A summary of generated features, their feature groups, and their dimensions is presented in Table 1.

In the process of filter feature selection, chi-2 test and F-test were used as filter methods, and the F-test showed a slightly better result. For the filter feature selection, we used the SelectKBest method, and the value of K was chosen to be 400 based on the better prediction performance (Figure 4).

**Figure 4.**
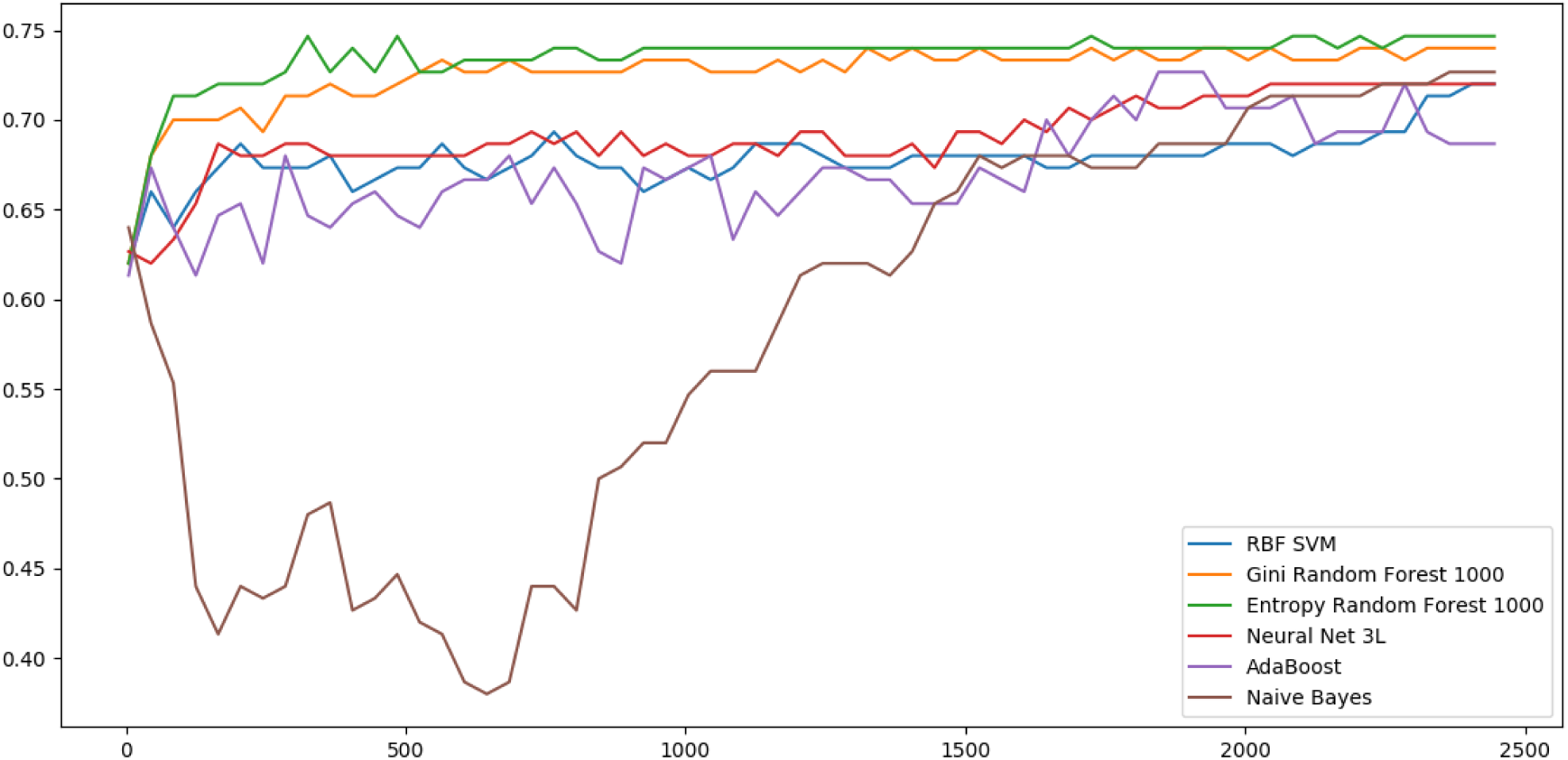
This illustrates the performance comparison of different algorithms for this classification task. The X-axis represents the number of features used for training (the K value in the SelectKBest method) and the Y-axis represents the demonstrated accuracy of each classifier during 6-Fold CV tests.

### 3.2. Model selection and evaluation

Figure 4 demonstrates the performance comparison of different classification methods with different numbers of features.

Our final model was chosen to be the RF, with information gain as the splitting criterion, due to better classification accuracy. Reported performance metrics in Table 2 were computed through 150 six-fold CV tests and ten holdout tests using the unseen test data (10% of the initial dataset, which was reserved at the beginning). In each CV and holdout test iteration, the samples were shuffled with different random seeds before splitting. Table 2 shows the comparison of TAXyl’s evaluation results in CV and holdout tests and Figure 5 illustrates the TAXyl’s performance through ten holdout tests on unseen data.

**Figure 5.**
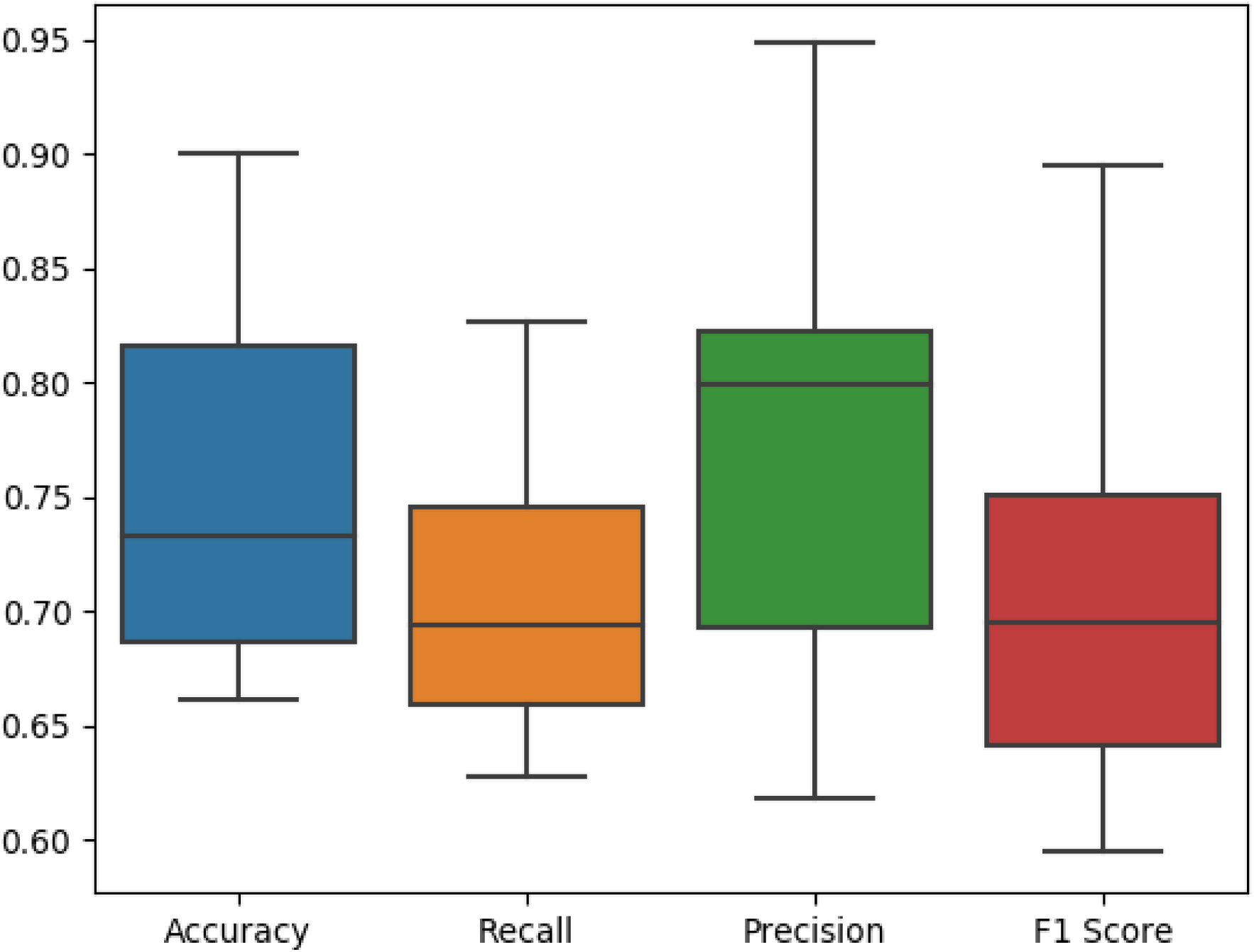
Box plot of TAXyl evaluation metrics during 10 times testing on holdout sets (unseen data).

### 3.3. Cloning, expression, purification, and characterization of xylanases with predicted thermal activity

Aiming at targeted enzyme isolation and screening of metagenomic data from sheep and cow rumen, TAXyl’s was implemented to obtain an *in-silico* estimation of the thermal activity status of numerous xylanase candidates prior to the cloning and other wet-lab experiments. Three probable xylanses from metagenomic sources with acquired thermal activity predictions were cloned, expressed, purified, and characterized. Three candidate enzymes produced from successful cloning, expression, and purification were named PersiXyn5, PersiXyn6 and PersiXyn7. These enzymes were subjected to further biochemical experiments. Furthermore, PersiXyn1 and PersiXyn2 from two of our previous studies were also used for purpose of validating TAXyl’s predictive ability. Table 3 contains the list of enzymes, primers and restrictions used in the cloning process, the prediction of TAXyl, and their experimentally identified optimum temperature.

### 3.4. Online Server

TAXyl is freely available at http://arimees.com. This web service is capable of receiving inputs in forms of FASTA file, amino acid sequence, or protein entry of xylanases from GH families 10 and 11 and returns their probable thermal activity status. TAXyl also enables users to export the selected features for their inserted protein sequences in CSV format (with the purpose of reproducibility of results or providing features for other machine learning tasks). This web-service is also accessible from the CBB lab website (http://cbb.ut.ac.ir), under databases and tools sub-menu.

## 4. Discussion

Xylanases are carbohydrate-active enzymes with multiple industrial applications and high commercial value (D. Kumar et al., 2017). Determining the optimum temperature of activity plays an important role in choosing the appropriate enzymes for specific purposes. Since the advent of next-generation sequencing and its accelerating improvements, getting access to metagenomic data is becoming increasingly easier and more affordable, and the only possible approach to analyze these extensive amounts of data is by fast and accurate computational methods.

One of the challenges of this study has been the limited number of xylanases with a reported optimum temperature in the literature and thus a smaller dataset. Therefore, we explored different machine learning methods to find the one with an acceptable interpretation of the data.

Our results indicated the competence of computational methods to address common problems in bioinformatics and the capability of sequence-based and length-independent protein descriptors for training supervised learning algorithms to effectively predict general enzymatic attributes (Pucci et al., 2014).

Most importantly, this methodology (Figure 3) can be generalized for other enzyme families and enzymatic properties. Through the implementation of this approach to develop various predictive models, the challenge of targeted enzyme identification as well as agile, effective, and inexpensive screening of high-throughput data can be addressed.

This tool helps to significantly reduce the number of potential candidate enzymes with a specific thermal activity profile prior to engaging the wet-lab experiments. Another potential usage for such tools in bioinformatics is in facilitating the engineering of enzymes through directed evolution to obtain biocatalysts with specific thermal dependence targeted at particular industrial purposes.

Unlike other studies designed to classify the status of enzymes’ thermal activity and stability (Gromiha & Suresh, 2008; Tang et al., 2017; Zhang, 2013) our model, extends the classification ability to the third class which are the hyper-thermophilic enzymes focusing on xylanase families. In comparison with our previous two studies, in which xylanase enzymes from the metagenomic source were identified and characterized by experimental techniques (Ariaeenejad, Hosseini, et al., 2019; Ariaeenejad, Maleki, et al., 2019), TAXyl enabled the identification of more putative thermophilic xylanases and enhanced the extended exploration of the metagenome. This tool was used to estimate the thermal activity of putative xylanases identified within the metagenomic samples and wet-lab experiments validated the predictions which were made previously.

## 5. Conclusion

In this study, we presented a novel method based on a supervised machine learning algorithm to predict the thermal activity of GH10 and GH11 xylanases. TAXyl uses sequence-based and length-independent protein descriptors to train a random forest classifier. TAXyl facilitated the targeted mining of three xylanases with specific thermal dependences from cow and sheep rumen metagenome.

The presented methodology for the development of this prediction tool can be generalized and implemented to model the functional properties and predict various attributes of a wide range of enzymes from different families. In case of availability of sufficient data, a possible direction for future works would undoubtedly be developing similar tools to predict the structural, functional, and thermodynamic properties of other enzyme families. The TAXyl web-service is available and provides users with a reasonably accurate prediction of any GH10 or GH11 xylanase thermal activity status.

## Supporting information

This supplemental document contains the dataset used for training of the predictive model.

## Supporting Information

Dataset which was used for this study. (.xlsx)

## Availability and implementation

**Prediction online webservice:** http://arimees.com/

**Codes:** https://github.com/mehdiforoozandeh/TAXyl

## Acknowledgments

This research was supported by a grant from the Agricultural Biotechnology Research Institute of Iran (ABRII), Institute of Biochemistry and Biophysics (IBB). Moreover, the authors declare that they have no competing interests.

## Abbreviations

AAC: Amino acid composition;
2AAC: Dipeptide composition;
3AAC: Tripeptide composition;
APAAC: amphiphilic pseudo amino acid composition;
CTD: Composition, Transition, Distribution;
CTF: conjoint triad features;
CV: Cross Validation;
GH: Glycoside Hydrolase;
MLP: Multi-Layer Perceptron;
PAAC: Pseudo amino acid composition;
QSO: Quasi-sequence Order;
RBF: Radial Basis Function;
RFE: Recursive Feature Elimination;
SOCN: Sequence order coupling number;
SVM: Support-Vector Machine

